# The Baraitser-Winter Cerebrofrontofacial Syndrome recurrent R196H variant in cytoplasmic β-actin impairs its cellular polymerization and stability

**DOI:** 10.1101/2025.09.12.675785

**Authors:** Éva Gráczer, Elena Battirossi, Tamás Bozó, Áron Gellért Altorjay, Katalin Pászty, Laura Harsányi, Johannes N. Greve, Irene Pertici, Massimo Reconditi, Nataliya Di Donato, Miklós Kellermayer, Pasquale Bianco, Andrea Varga

## Abstract

Variants in cytoskeletal actin encoding genes are associated with a broad spectrum of disorders, called non-muscle actinopathies. Among them, Baraitser-Winter cerebrofrontofacial syndrome (BWCFF) displays the most severe symptoms, such as intellectual disability and epilepsy. The exact consequences of the mutation on actin’s properties, however, are not fully understood. Here we explored the cellular effects of the R196H mutation in patient-derived fibroblasts. We show that the heterozygous mutation causes an actin polymerization defect in cells, leading to a fifty percent decrease in filamentous (F-) actin content. This effect can be rescued by the addition of the actin-polymerizing and stabilizing drug, jasplakinolide. We observed no significant defects either in the organization of the cellular actin cytoskeleton, analyzed by superresolution (STED) microscopy, or in the structure of purified filaments stabilized with phalloidin, explored with atomic force microscopy (AFM). The reduced F-actin content correlated with an approximately fourfold reduction in the stiffness of patient-derived cells probed with AFM. Manipulating the cells by mechanical forces through the application of the Dual Laser Optical Tweezers (DLOT) technique suggests that the mutation weakens the attachment of cytoskeletal actin to the plasma membrane. Inducing dynamical reorganization of actin by uniaxial stretching revealed that the interaction of cofilin with actin is also weakened by the mutation. Based on the existing cofilin-actin structures, the binding of cofilin may weaken the interaction of the neighboring residue E195 with K113, one of the lateral contacts stabilizing the filament. Thus, the mutation possibly exerts its effect through the destabilization of the interfilament interactions, potentially interfering allosterically with cofilin binding during actin depolymerization.

## INTRODUCTION

Actin is one of the most highly conserved proteins and plays crucial roles in numerous cellular functions and processes, such as cell division, contraction, adhesion and migration. Actin forms filaments within the cells, which provide structural support and help the cell to maintain its shape and internal organization or develop force (1). To achieve its versatile functions, actin assembly and disassembly are spatially and temporally regulated by robust mechanisms with a large set of actin-binding proteins (ABPs) (2).

Humans express six actin isoforms (3). Four of them are muscle-type actins (α-actin isoforms of skeletal muscle, smooth muscle, cardiac muscle and γ-actin isoform of smooth muscle cells encoded by *ACTA1*, *ACTA2*, *ACTC1* and *ACTG2*, respectively), and two are cytoplasmic forms (β- and γ-actin isoforms, encoded by the *ACTB* and *ACTG1* genes, respectively). The actin isoforms do not differ by more than 7% at the primary amino acid sequence level. The two most similar actin isoforms, β-cytoplasmic (β-actin) and γ-cytoplasmic actin (γ-actin), differ by only four amino acids, which are clustered at the N-terminus of the 375AA-long protein. Despite their nearly identical sequence, the cytoskeletal actin isoforms display different polymerization properties, are localized in different parts of the cell, display preferred interactions with different ABPs and support different functions (4–7). In fibroblasts and epithelial cells, β-actin is preferentially localized in stress fibers and circular bundles at cell-cell contacts, suggesting its role in cell attachment and contraction (8). In moving cells, γ-actin is mainly organized as a meshwork of cortical and lamellipodial structures, suggesting a role in cell motility. In resting cells, γ-actin is also recruited into stress fibers. Any change that affects actin dynamics and monomer-polymer balance has a dramatic impact on the cell. *De novo* mutations in all six genes encoding actin isoforms can cause human diseases classified into muscle and non-muscle actinopathies (NMAs). Both types are autosomal-dominant disorders with many mutations resulting in heterozygous missense changes. This suggests that the mutant actin monomers are produced and incorporated into actin filaments responsible for the phenotypic changes (9).

To date, a broad spectrum of diseases, referred to as NMAs, associated with mutations in the *ACTB* and *ACTG1* genes encoding the cytoskeletal β-actin and γ-actin isoforms, respectively, has been identified. These include a range of rare syndromes that are often associated with brain malformation (7, 10–14). Despite the intense research in the field, the available neuropathological data are very limited (15–17). Moreover, the impact of allosteric perturbations such as disease-causing mutations on the structure and function of discrete actin-based complexes and their various effects on cell morphology, behavior, morphogenesis, and development are unknown.

Baraitser-Winter cerebrofrontofacial syndrome (BWCFF) is a subtype of NMAs associated with typical craniofacial features and cortical malformations, leading to intellectual disability. Several β- and γ-actin mutations are associated with BWCFF, such as G74S, T120I and R196H or R196C of β-actin and S155F and T203M of γ-actin. Among them, R196 is a mutation hot spot, which is frequently observed in BWCFF patients, especially the variant R196H (14). There are, however, other mutations associated with less severe symptoms, such as G302A, S368 frameshift and S338-I341 deletion in β-actin (13, 18). There is an attempt to categorize these mutations according to the properties of actin affected, such as haploinsufficiency (actin monomer quantity insufficient), polymerization-depolymerization efficiency, binding ability to different ABPs, or the assembly of toxic oligomers not compatible with normal cell functions (19). To date, only a few mutations have been investigated *in vitro* by analyzing the properties of the recombinant proteins with the specific mutations such as E334Q (of γ-actin) and S368 frameshift variant (of β-actin) (19, 20). The mutant E334Q affected actin binding to cofilin and to myosin motors, while the S368 variant showed a defect in actin polymerization and profilin binding. The same researchers investigated the effect of R196H mutations and found that it changed the rate of actin polymerization and depolymerization and decreased the Arp2/3-mediated *in vitro* filament branching (21).

Residue R196 of β-actin is located close to the monomer-monomer interface, but it is not involved directly in the stabilization of the filament through interstrand contacts (Figure 1A). By contrast, the neighboring E195 forms a salt bridge with K113 from the adjacent monomer (Figure 1B and C). This interaction is part of an allosteric route, which might be involved in propagating structural changes to the neighboring filament (22). Residue K113 is located close to the backdoor of the P_i_-release route, which is predominantly closed in the core of the filament (due to steric hindrance, partly by the interaction of residues R177 and N111, Figure 1B) (23). The question arises whether the mutation might affect the binding of cofilin to actin, which is related to the release of P_i_. Cofilin has a preference for binding to the ADP-bound form of actin, and it binds on the opposite side of the interfilament interface by bridging two actin monomers (Figure 1A).

**Figure 1.**
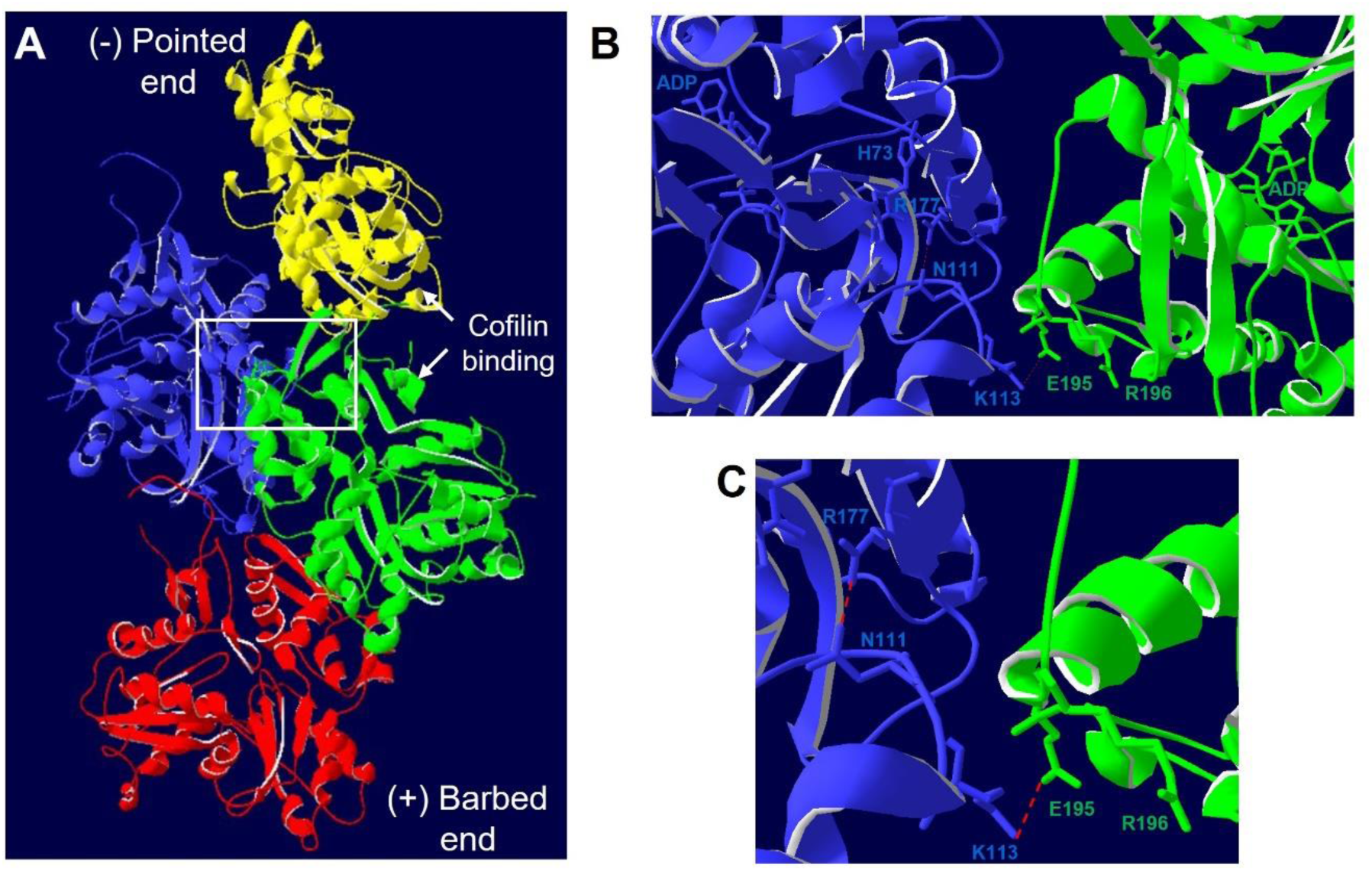
Location of the R196H residue in the F-actin structure **(A)** Structure of cytoskeletal β-actin (PDB ID: 8OI6, (23)), protomers are coloured as yellow, blue, green and red from the pointed (-) end to the barbed end (+). **(B)** The filament interface between the blue and green protomers is enlarged and rotated to highlight the ADP binding pocket and the backdoor residues (blue protomer) important for Pi release, which are shown in stick models together with the residue R196, its neighboring residue, E195 (green protomer) and the residue K113 (blue protomer). The interaction between the latter residues is one of the lateral contacts responsible for filament stabilization (66). **(C)** A further zoomed-in view of the interfilament interface showing the contacts between R177 and N111 (blue protomer), which is important to slow down Pi release in the core of the filament as well as the lateral contact between K113 (blue protomer) and E195 (green protomer).

Reorganization properties of the actin cytoskeleton can be studied by applying mechanical forces on single cells or cell monolayers. The actin cytoskeletal response to mechanical forces and the role of specific actin mutations therein can be studied in different experimental settings. During cell stretching, the involvement of the ABPs cofilin and the formin Diaphanous were identified in the reorganization of the actin cytoskeleton (24). In addition, myosin II “polarization”, i.e., reorganization to the stretched junctions was also observed and its extent was proportional to the applied strain (and stress). We have found previously that, in response to uniaxial stretch, the extent of cofilin reorganization from the cell periphery correlated with the strength of cell-cell junctions in endothelial cells (25), showing the importance of actin in cell-cell communication. The mechanical perturbation of live cells can be achieved on different force and directional scales by atomic force microscopy (AFM). Cell stiffness, determined by AFM, might be modulated by all three major components of the cytoskeleton: actin cytoskeleton, microtubules and intermediate filaments. However, cell stiffness might be dominated mostly by the actin cytoskeleton (i.e., the amount of stress fibers (26)), but changes in the microtubular structure have also been shown to modulate cell stiffness (27). The plasma membrane of animal cells, which conforms to the underlying cytoskeleton, exhibits dynamic interactions that may be probed by methods, such as optical tweezers, which enable manipulation in the low-force regime. One key application is the study of membrane tethers (or nanotubes), which can be spontaneously generated by cells or artificially extracted by applying local forces (28). These tethers are critical in processes like filopodial extension, endocytosis, and the formation of nanotubular connections between cells (29). Artificially generating membrane tethers provide valuable information about membrane mechanics. Another key optical tweezers application in cell mechanics is nanoindentation, which measures cell deformation under an external pushing force (30, 31). Dual Laser Optical Tweezers (DLOT) offer a powerful advantage over single-beam optical tweezers owing to the higher forces while maintaining precision (32). DLOT enables more effective probing of the mechanical properties and dynamics of the membrane-cytoskeleton system, which is essential for understanding cellular mechanics at a deeper level.

Here, we investigated how the single point mutation R196H in β-actin affects the organization of the actin filaments in patient-derived fibroblasts and the reorganization of actin upon mechanical challenge by monolayer stretching. Furthermore, the effect of the mutation on the mechanical properties of the recombinant actin as well as the fibroblasts were measured by AFM, reflecting changes in overall cellular rigidity. In parallel, DLOT was employed to specifically probe the mechanical response of the plasma membrane–cortical actomyosin system. The patient-derived fibroblasts were used as a relevant model to evaluate whether the mutation could modify any aspects of actin organization at the cellular level. We show that the mutation reduced the amount of filamentous (F-)actin in the cells, which was also reflected in the decreased cellular stiffness. We also identified that the mutation affects the orientation of the actin filament bundles by perturbing Arp2/3 binding, as well as actin reorganization by disturbing cofilin binding.

## RESULTS

### R196H mutation impairs actin filament stability, organization and cell stiffness in patient-derived fibroblasts

In the present work, we investigated the structural, dynamic and mechanical consequences of the BWCFF-associated mutation R196H in cytoskeletal β-actin. To find out whether the polymerization defect of the actin variant R196H, which has been observed *in vitro* (21), might be attenuated in cells by the presence of the wild type allele, we analyzed the F-*versus* total actin ratio in patient-derived fibroblasts. To investigate whether the mutation influences the amount of the actin filaments, we lysed wild type and R196H mutant patient-derived fibroblast cells and separated F-actin from G-actin by ultracentrifugation. We found that the amount of F-actin decreased by 50% upon the mutation (Figures 2A and B), as analyzed with either a β-actin or a pan-actin antibody. To explore whether the polymeric form of actin could be recovered, we treated the cells with jasplakinolide. Upon adding jasplakinolide, the amount of F-actin increased two-fold in both the wild type and mutant cells. Thus, the effect of the R196H mutation can be rescued with jasplakinolide. To explore whether the mutation decreases the total amount of β-actin in the cells, we analyzed their lysate by western blotting (Figure 2C). There was no significant difference in the amounts of β-actin and house-keeping proteins (GAPDH and tubulin) between the mutant *versus* wild type cells. Thus, the mutation does not affect the total amount of actin, but it decreases that of F-actin.

**Figure 2.**
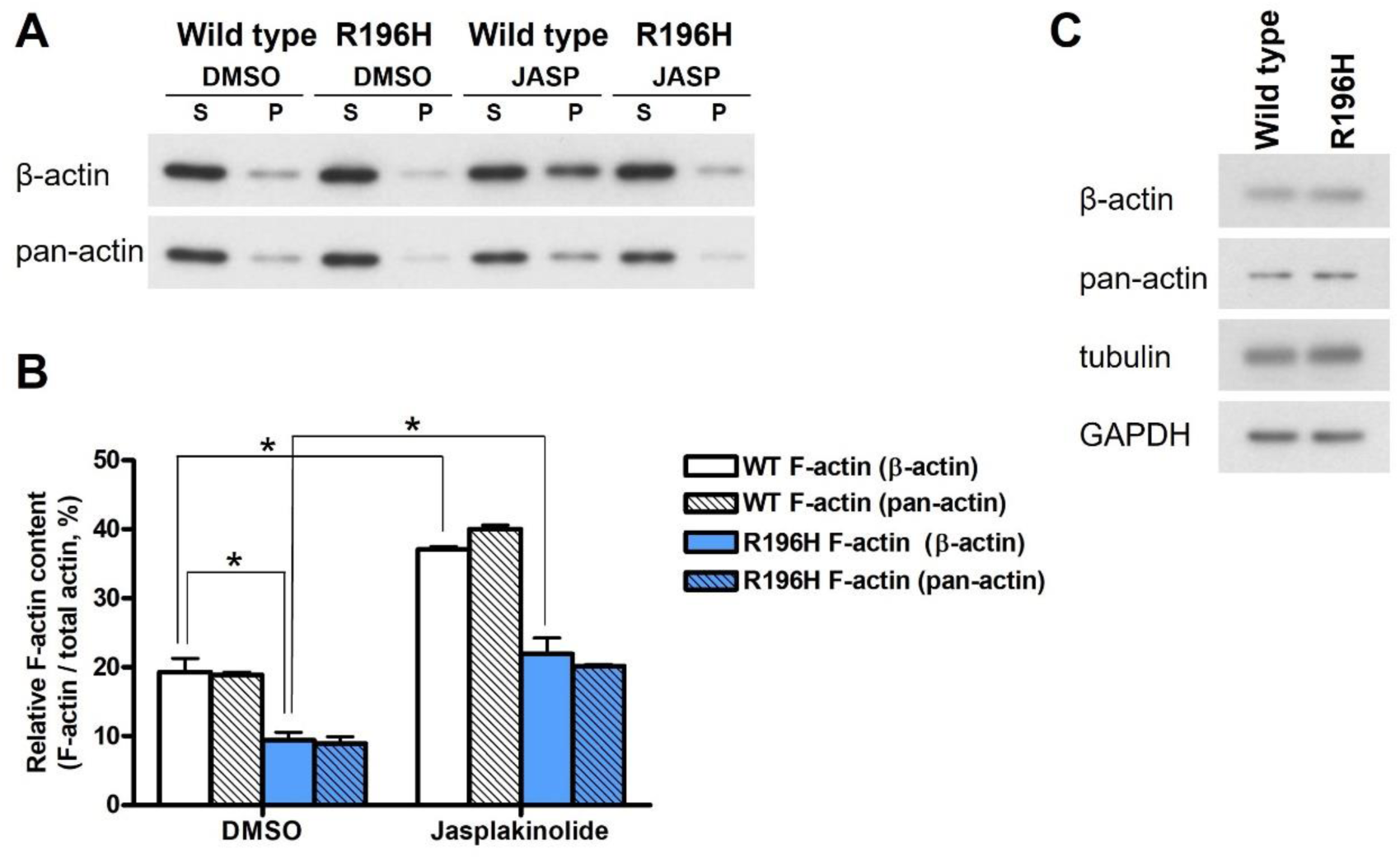
R196H mutation impairs actin filament stability in patient-derived fibroblasts **(A)** F-actin (recovered from the pellet after ultracentrifugation, P) and G-actin (supernatant, S) content of DMSO control or jasplakinolide-treated (0.1 μM, 30 min) wild type (WT) or R196H mutant fibroblast cells were determined by western blotting using either β-actin or pan-actin antibodies. **(B)** The amount of F-actin was quantified for each condition as the ratio of F-actin and the total actin (the sum of F- and G-actin) for each sample separately. **(C)** Total lysates of wild type and R196H mutant fibroblasts were immunoblotted and labeled with the antibodies of β- and pan-actin. Tubulin and GAPDH were also labeled as loading controls. Results of two independent experiments are shown. Data were analyzed by two-way ANOVA, followed by Bonferroni’s multiple comparison test expressed as mean ± SD (*p < 0.05).

To find out whether the decreased amount of F-actin is due to the decreased filament bundle width (i.e., fewer filaments in the bundle), or rather the organization of F-actin is different in mutant cells, we fixed the cells and labeled them with a phalloidin derivative compatible with STED imaging (Figure 3A). In wild type cells, the mean value of the width of actin filament bundles (fitted from a Gaussian distribution, Figure 3B, Supplementary Figure 1A) was 0.136 ± 0.002 µm. By taking the published value for the width of the double helix (7-9 nm, (33)), we can estimate that about 15-20 filaments are present in the actin bundles. We also pre-treated the cells with jasplakinolide to monitor its effect on filament width. Based on the Gaussian distribution of the quantified fluorescence signals (Figure 3B, Supplementary Figure 1A-C) we could not see any significant difference in filament width upon either the mutation or jasplakinolide treatment. To analyze the organization of the actin filament bundles we used live cell confocal imaging (Figure 3C). To quantify the degree of alignment of actin filament bundles, a directionality analysis was carried out. A comparison of the Gaussian fit of the directionality histograms of wild type and R196H mutant cells is shown in Figure 3D. From the analysis of such types of histograms (Supplementary Figure 1D and E), we have found a difference in the dispersion of the orientation of actin filament bundles of wild type and mutant cells without jasplakinolide treatment. Interestingly, jasplakinolide treatment of wild type cells significantly reduced the dispersion to the level indistinguishable from that of the R196H cells (Figure 3E). These results imply that actin filaments within the bundle are more aligned in the mutant cells, suggesting that jasplakinolide might regulate the alignment of the actin fibers *in vitro*. To uncover whether the orientation of the bundles is regulated by the Arp2/3 complex, we treated wild type cells with the Arp2/3-specific inhibitor CK-666 (Figure 3F). We found that the dispersion of the orientation of actin bundles decreased upon CK-666 treatment and its value became very similar to the value obtained with the R196H mutant cells (Figure 3G and Supplementary Figure 1F and G).

**Figure 3.**
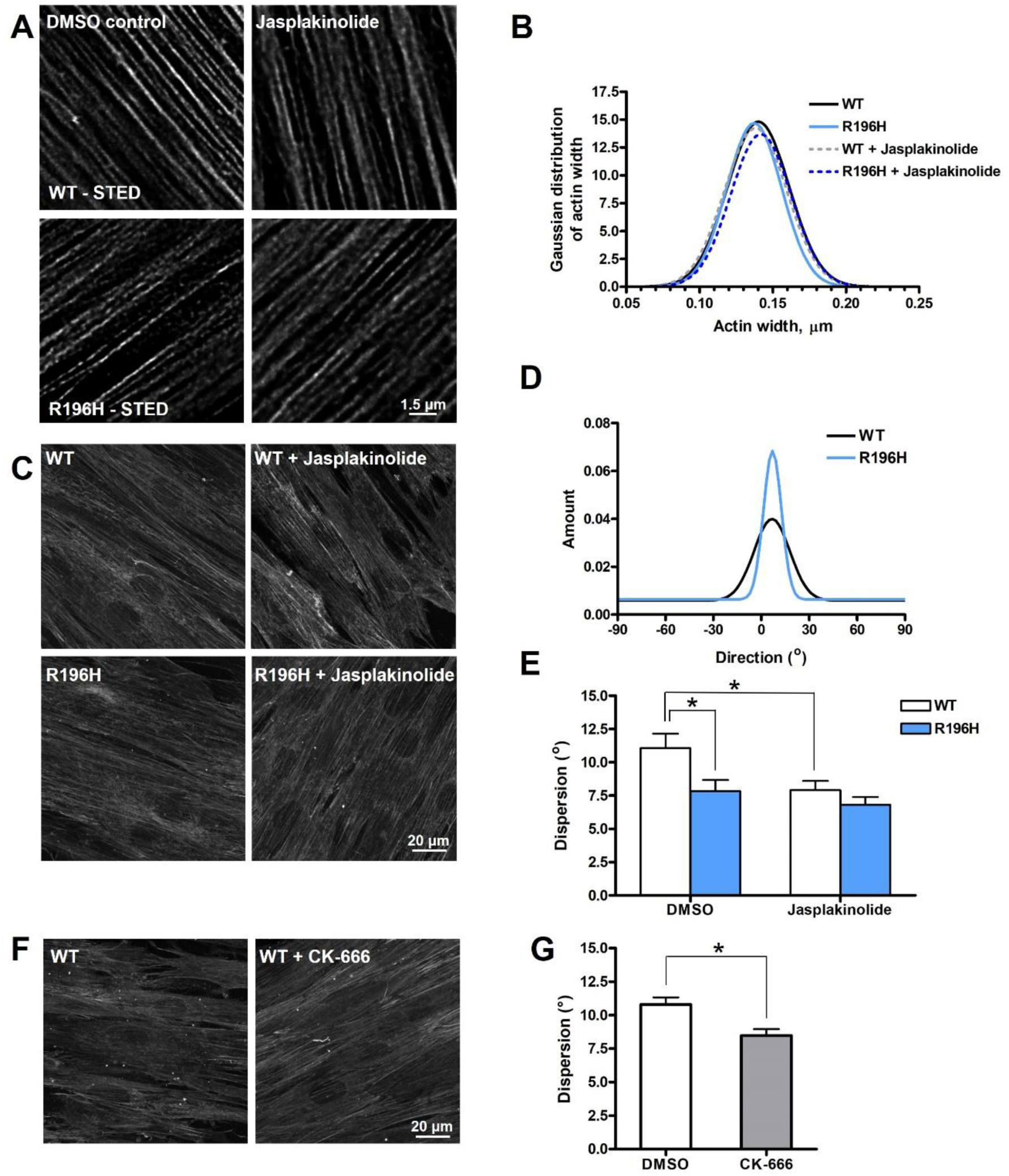
R196H mutation impairs actin filament organization in patient-derived fibroblasts **(A)** Superresolution STED images of DMSO control or jasplakinolide-treated (0.1 μM, 30 min) wild type (WT) or R196H cells. **(B)** Gaussian fit of the distribution profile of actin width (Supplementary Figure 1A-C) quantified from the line profile after image deconvolution. **(C)** Confocal images of live cell actin staining of DMSO control or jasplakinolide-treated (0.1 μM, 30 min) WT or R196H cells. **(D)** Directionality analysis showing an example of the Gaussian distribution of filament orientations of phalloidin-stained WT or R196H cells. **(E)** Dispersion calculated from the width of the Gaussian distribution determined from the phalloidin-stained confocal images of WT or R196H cells. **(F)** Confocal images of live cell actin staining of DMSO control or CK-666-treated (100 μM, 60 min) WT cells. **(G)** Dispersion calculated from the width of the Gaussian distribution determined from the phalloidin-stained confocal images of WT DMSO control or CK-666-treated cells. Data are from two independent experiments. Data were analyzed by two-way ANOVA, followed by Bonferroni’s multiple comparison test expressed as mean ± SD (*p < 0.05).

To investigate whether the mutation-associated changes in the F-actin content and/or filament directionality change the stiffness of the cells (Figure 4A), we carried out atomic force microscopy (AFM) measurements. Representative force curves and Young modulus distributions of wild type and R196H cells are shown for comparison in Figure 4B and C, respectively. Remarkably, the R196H actin mutation led to a fourfold reduction of the mean elastic modulus and to a fivefold decrease in its distribution (measured as the full width at half maximum, FWHM) (Figure 4C). Thus, the cells carrying the mutation are significantly more compliant than the wild type ones. To analyze the effects of the mutation on the structure of individual actin filaments, wild type and R196H recombinant actin purified from baculovirus-infected insect cells were polymerized in the presence of phalloidin, and the topography of the filaments was analyzed with AFM. The actin filament samples were added to a supported lipid bilayer system, on which actin paracrystals formed, enabling us to measure not only the topographical structural details of individual filaments with high resolution, but their interactions and organization as well (Figure 4E). Even in the presence of phalloidin, fewer actin filaments were formed by the mutant actin. The quantitative analysis of the monomer-monomer distance within a single actin filament (Figure 4F) as well as the half-helical pitch (Figure 4G), did not show differences upon the mutation, and the obtained mean values were close to the published values (34) of 5.5 nm and 36 nm for the monomer-monomer distance and for the half-helical pitch, respectively.

**Figure 4.**
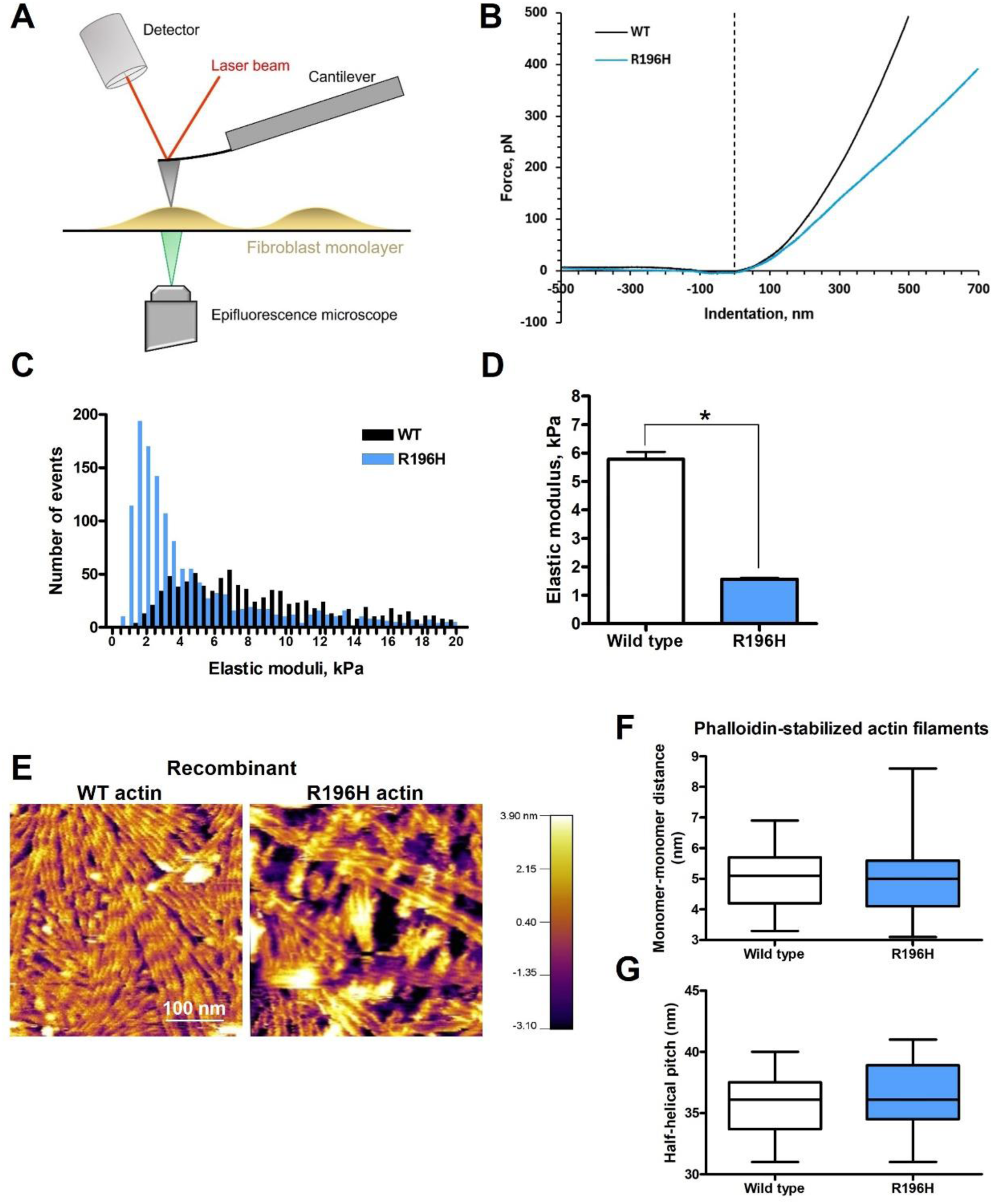
R196H mutation decreases the elastic modulus of patient-derived fibroblasts **(A)** Schematic representation of the experimental set-up of atomic force microscopy measurements carried out with wild type (WT) and R196H mutant live fibroblasts. **(B)** Representative force-indentation curves of WT and R196H fibroblasts. **(C)** Distribution of the elastic moduli of wild type (black) and R196H (blue) calculated from the force-distance curves as described in the Materials and Methods section. **(D)** Mean values of the Gaussian fit of the histograms (calculated from the log-normal distribution) are shown for wild type (white) and R196H (blue) cells. **(E)** Topography of recombinant wild type or R196H actin paracrystals determined by atomic force microscopy. Monomer-monomer distance **(F)** as well as half-helical pitch **(G)** were determined and plotted for wild type and R196H recombinant actin. Data are from two independent experiments. Data were analyzed by one-way ANOVA, followed by Bonferroni’s multiple comparison test expressed as mean ± SD (*p < 0.05).

### R196H mutation affects the attachment of actin to the plasma membrane

Using the DLOT system with a micro-bead probe, we applied forces to the beads in the physiologically relevant ten-piconewton range to induce measurable cell membrane indentation and extract membrane tethers by pulling the plasma membrane. A polystyrene bead was optically trapped near a cell (Figure 5A). The cell was then moved toward the bead to establish contact and induce lateral indentation (35). Figure 5B shows the force response (blue trace) to ramp-and-hold push-and-pull protocols under position feedback (red trace) related to the plasma membrane’s lateral-indentation followed by tether extraction. Importantly, the force values reached during tether elongation were highly consistent across cycles, strongly suggesting that single tether extraction events were reliably isolated and used for analysis. Upon retraction, membrane tethers formed with lengths between 10-50 μm, and tether formation persisted through subsequent cycles without the need to repeat the indentation.

**Figure 5.**
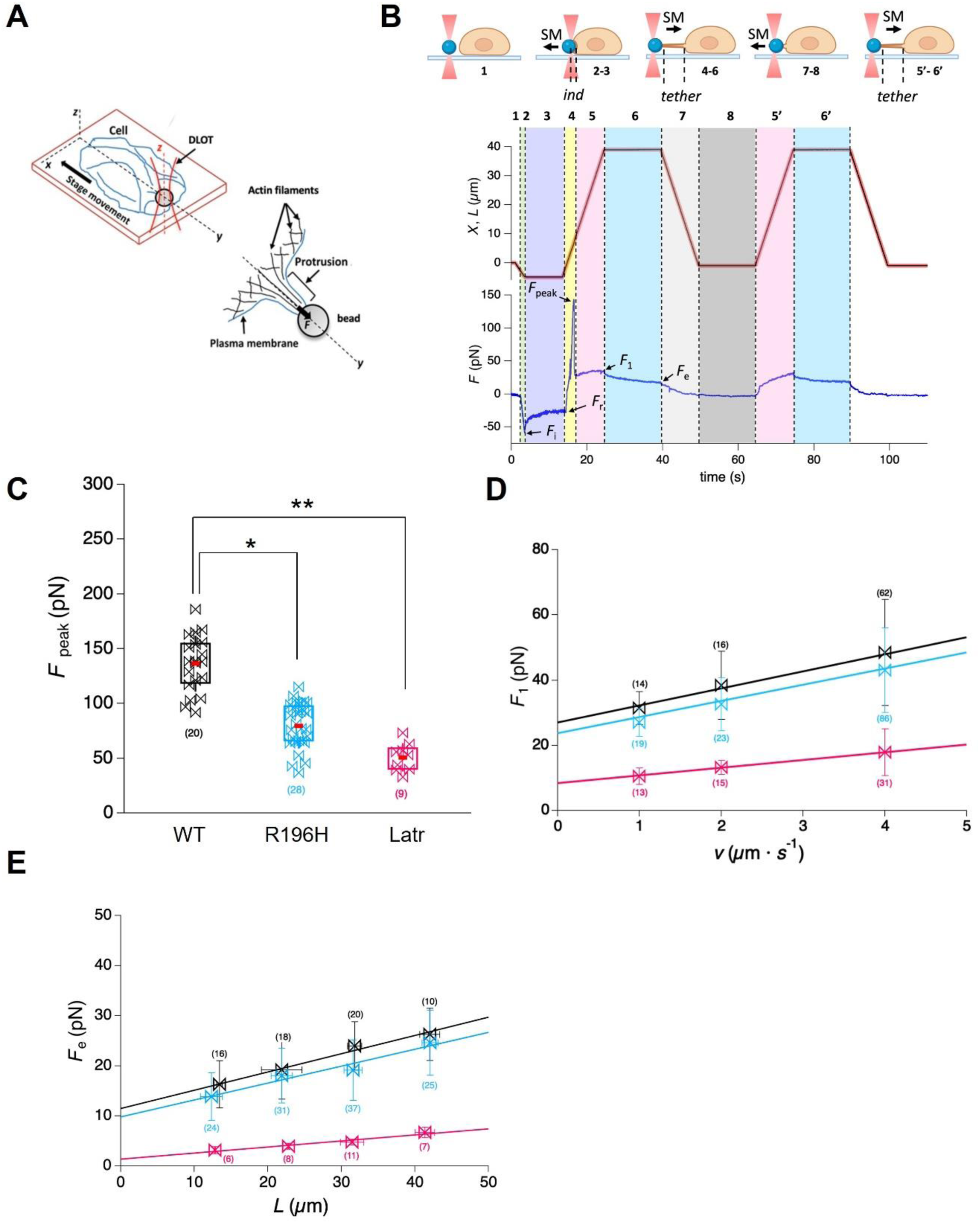
R196H mutation affects the attachment of actin to the plasma membrane **(A)** Schematic representation of the protrusion induced on an adherent living cell using a laser trapped bead. Mechanical stimuli under position feedback are applied locally on the cell membrane with the laser-trap polystyrene bead (3 µm diameter) functionalized with -NH2 groups to increase the probability of interaction with the cell membrane. A protrusion is formed in response to a pulling force (F) applied on the cell membrane along the y direction. **(B)** Top Panel: Schematic representation of the mechanical protocol applied to a cell. Specific phases of the protocol are numbered as detailed in the text. Bottom Panel: The piezo position (red trace), membrane position (black trace), and force response (blue trace, negative resisting to compression, positive resisting to extension) plotted as a function of time. The colors help identifying the phases: [1] white, [2] light green, [3] light purple, [4] yellow, [5 and 5’] pink, [6 and 6’] light blue, [7] light gray, [8] gray. The key forces are defined as follows: *F_i_*, the force value attained at the end of the indentation ramp; *F_r_*, the force attained after the indentation relaxation process; *F_peak_*, the force peak resisting to extension; *F_1_*, the force reached during tether elongation; *F_e_*, the equilibrium value after tether relaxation. SM refers to the movement of the stage. **(C)-(E)** Color code: WT (black), R196H (light blue), Latr (pink). The number of experiments carried out for each condition is in brackets. Data were analyzed by two-way ANOVA, followed by Bonferroni’s multiple comparison test (*p < 0.05, **p < 0.01). Data are presented as mean ± SD. **(C)** Box plot showing *F_peak_* values (pN) for different experimental conditions. The box boundaries are defined according to Minitab criteria, where data points within one standard deviation of the mean are considered within the expected range. Given that the distribution of data points is not Gaussian, no outliers were removed. The definition of the error by one standard deviation ensures a robust depiction of variability that reflects the non-Gaussian nature of the dataset. The red lines represent the mean value for each boxplot. **(D)** Linear dependence of *F*_1_ on *v* for elongations greater than 35 μm. The slope of the linear fit provides an estimate of the effective viscosity (γ_eff_), which is 1.05± 0.45, 0.88± 0.35, and 0.24 ± 0.11 pN·s/μm for wild type, R196H, and latrunculin A–treated cells, respectively. The y-intercept (27.4 ± 3.7, 23.7 ± 4.2, and 7.9 ± 2.4 pN for wild type, R196H, and latrunculin A–treated cells, respectively) provides an estimate of *F*ₑ for 38 μm < L < 43 μm, which is in good agreement with the value obtained from the *F*ₑ–*L* relation in panel E. **(E)** Linear dependence of *F_e_* on the tether length *(L)*. The panel illustrates the equilibrium force (*F*_e_) at various tether elongation lengths (*L*) for each experimental condition.

During ramp-shaped push, when applied over a short duration, both force and indentation depth increased linearly up to a value attained at the end of the ramp, characterized by a force *F_i_* and an indentation *d_i_*, suggesting an elastic response (Figure 5B, phase 1 and 2, and Supplementary Figure 2A). Following the end of the ramp (Figure 5B, phase 3), force relaxation occurred, which implies the return of the bead toward the trap center with further movement of the membrane and corresponding to further increase in indentation. This phase reveals the viscoelastic nature of the response to indentation, which reaches a near-equilibrium state characterized by a force *F_r_* and an indentation *d_r_*. From the values *F_i_* and *F_r_* two apparent elastic moduli can be estimated: the “instantaneous” elastic modulus *E_i_*, given by the parallel of the three springs (*k_0_+k_1_+k_2_*), and the “static” elastic modulus *E_r_*, due to the only element *k_0_* (for further details see the Materials and Methods section, Eq. 4, Supplementary Figure 2B). While there were no significant differences in the mechanical properties between wild type (*E_i_*=57.7±19.8 Pa and *E_r_*=21.9±7.9 Pa) and the R196H fibroblast (*E_i_*=71.3±28.3 Pa and *E_r_*=21.3±7.4 Pa) cell lines, wild type fibroblasts treated with latrunculin A showed a significant reduction in stiffness (*E*_i_ of 10.2 ± 5.1 Pa and *E*_r_ of 5.3 ± 2.1 Pa), demonstrating that disruption of the actin cytoskeleton leads to decreased mechanical integrity (Supplementary Table 1).

Tether formation revealed a characteristic force peak (*F_peak_*) (Figure 5B, phase 4), which is the initial force required to detach the membrane from the cell cortex, and it is related to the strength of membrane – cell cortex adhesion. The *F_peak_* value of the wild type cells (138±25 pN) was 1.75-times larger than in the R196H mutant cells (83±20 pN) (Figure 5C). Depolymerization of actin by latrunculin A in wild type fibroblasts reduced the value of *F_peak_* further (51±12 pN) (Supplementary Table 1). These results imply that the mutation affects the coupling of the plasma membrane and the actin cytoskeleton of the cell cortex.

During tether elongation (Figure 5B, phase 5), force increased and reached *F_1_* at the end of the ramp, then during the hold phase it relaxed bi-exponentially to *F_e_* (Figure 5B, phase 6). From the linear relation of *F*_1_ versus velocity (*v*) (Figure 5D) we obtained the tether’s effective surface viscosity (γ_eff_). Our results indicate that this parameter does not differ significantly for wild type (γ_eff_=1.05 ± 0.45 pN·s/µm) and the mutant cell line (γ_eff_=0.88 ± 0.35 pN·s/µm). Latrunculin A treatment of wild type cells (γ_eff_=0.24 ± 0.11 pN·s/µm) confirmed the contribution of the actin cytoskeleton to *F*_1._ The equilibrium force (*F*_e_) was measured at the end of tether elongation across different pulling lengths (*L*). From their linear dependence (Figure 5E), the tether stiffness *k_0_* (defined as the slope of the *F_e_*-*L* relationship) and the tension necessary for generating and maintaining the membrane bending at the base of the tether, *F_base_,* (the intercept) can be derived. The mutation did not have a significant difference either on the *k_0_* (0.36±0.05 pN/μm for wild type, and 0.34±0.05 pN/μm for the R196H mutant) or on the *F_base_* (11.5±1.8 pN for wild type, and 9.8±1.4 pN for the R196H mutant) value. Lantrunculin A treatment of wild type cells decreased both the *k_0_* and the *F_base_* value (*k_0_* = 0.13±0.02 pN/μm and *F_base_* =1.4±0.4 pN), which indicates the role of actin filaments in the force relaxation.

Tether radius (*r_t_*) was not changed upon mutation (108±6 nm for wild type, and 117±12 nm for the R196H mutant cells) but increased significantly upon latrunculin A treatment (184±6 nm), indicating reduced cytoskeletal support. The evaluation of *F_peak_* in relation to the equilibrium force (*F_e_*) (calculated by taking into account the fact that a circular contact was formed between the bead and the cell membrane) led us to the normalization of *F_peak_* values (named as *F*_peak_*, see Materials and Methods section for the details). The *F*_peak_* values showed a reduction upon the R196H mutation (and also upon latrunculin A treatment), giving a value of 49±7 pN/nm^2^ for wild type, 32±6 pN/nm^2^ for the R196H mutant and 0.87±0.20 pN/nm^2^ for wild type cells, treated with latrunculin A, highlighting the role of the actin filaments in resisting local stress (Supplementary Table 1).

Finally, membrane tension (*T*) and bending modulus (*e*_b_) were calculated from *r_t_* and *F_e_*. Our results show no significant difference between wild type (*T*=17±2 μN/m and *e_b_*=456±83 zJ) and the R196H mutant (*T*=14±2 μN/m and *e_b_*=303±77 zJ) fibroblasts. Lantrunculin A treatment of wild type cells, indeed, decreased both the membrane tension (1.0±0.2 μN/m) and the bending modulus (104±15zJ), confirming that actin disruption compromises mechanical resilience.

### R196H mutation increases the dynamic reorganization of actin by enhancing cofilin dissociation

To determine whether the mutation affects the reorganization of actin, wild type and R196H fibroblasts were exposed to uniaxial stretch. The cell monolayers were kept either in a non-stretched state, or they were stretched by 30% of their original length. The cells were kept stretched for 15 minutes, fixed and stained for actin and cofilin (Figure 6A-H). Comparison of phalloidin staining of stretched wild type and R196H mutant cells showed that the actin bundles formed in the periphery of mutant cells in the direction of stretch are thinner compared to that of wild type cells (Figure 6B and D). In accordance with our qualitative understanding of cofilin reorganization upon stretch in endothelial cells (25), cofilin, indeed, was reorganized from the cell periphery upon stretch (data are shown in Figure 6I) both in wild type and in mutant cells. The relative movement of cofilin from the cell periphery increased in the presence of the mutation. The finding indicates that cofilin dissociation from peripheral actin upon stretch is enhanced in the presence of the R196H mutation. Since the binding site of cofilin is far from the site of the mutation, cofilin binding might be allosterically perturbed upon the mutation (Figure 6J, K).

**Figure 6.**
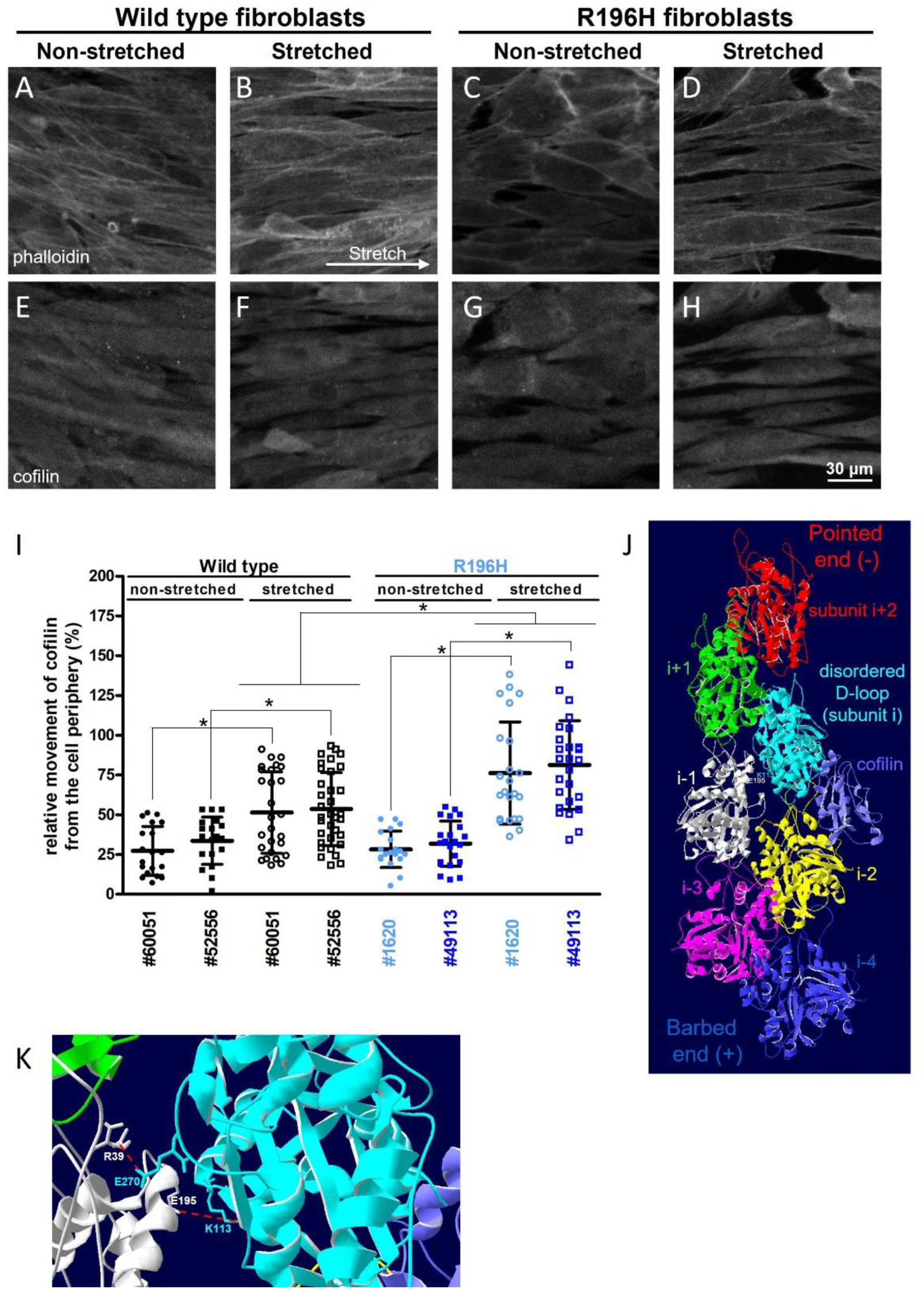
R196H mutation increases the dynamic reorganization of actin by enhancing cofilin dissociation **(A-H)** Immunofluorescence analysis of wild type (A, B, E, F) or R196H (C, D, G, H) fibroblasts kept either non-stretched (A, C, E, G) or stretched (B, F, D, H) for 15 minutes. Samples were stained with either phalloidin (A-D) or for cofilin (E-H). **(I)** Quantification of the relative movement of cofilin from the cell periphery. Non-stretched samples of wild type cells, isolated from two patients (#60051 and #62556), are shown in black, filled circles/squares, while stretched samples are represented by black open circles/squares. Non-stretched samples of R196H cells (#1620 and #49113) are shown in blue, filled circles/squares, while stretched samples are represented by blue open circles/squares. The rationale of the quantification of the relative movement of cofilin is shown in Supplementary Figure 3A-H. Data are from three independent experiments. Data were analyzed by one-way ANOVA, followed by Bonferroni’s multiple comparison test expressed as mean ± SD (*p < 0.05). **(J)** Cofilin (purple) binding to the actin filament as represented by the published cryo-EM structure (PDB code: 6UBY, (42)). The actin protomers within the filament are colored differently and their relative position to cofilin is labeled as described in (42). The flexibility of the D-loop is increased in the cyan protomer (in the one closer to the pointed end of actin). The position of the interfilament interaction between E195 (located in the white actin protomer) and K113 (located in the cyan protomer) is also labeled. **(K)** Zoomed-in view of the interfilament interactions between the white and the cyan actin protomers showing the location of the salt bridges E195 (white)-K113 (cyan) and R39 (white)-E270 (cyan).

## DISCUSSION

Our study demonstrates that one of the hot-spot mutations of BWCFF, R196H contributes to the instability of the actin filaments, even in the presence of the wild type allele. Our results also show that although the overall amount of β-actin remains unchanged in mutant cells, the ratio of filamentous (F-) to monomeric (G-) actin is significantly decreased, suggesting a selective impairment of filament stability rather than expression. This finding indicates that the R196H mutation destabilizes actin filaments by interfering with filament assembly or promoting depolymerization, without affecting actin synthesis. Indeed, the recombinant actin having the same mutation has been shown to have a defect both in filament assembly and depolymerization (21). To “amplify” any conformational problem of the mutant recombinant actin, we investigated the structure of individual actin filaments within an actin paracrystal (36), where a side-by-side array of filaments are formed on a positively charged lipid surface. The structure of the phalloidin-stabilized mutant filament could not be distinguished from its wild type counterpart, although fewer filaments were formed from the mutant recombinant actin. These results point out that the mutant actin might be incorporated into the actin filaments also in the cells, but due to its instability (increased depolymerization rate) the total amount of F-actin is decreased compared to the “pure” wild type case. In the cells, the actin filament bundles have very similar widths and there is a difference only in the orientation of the filaments. Mutant filaments run more parallel with each other, and this can be explained by the observed weakening of the interaction of recombinant actin with the Arp2/3 complex (21). Indeed, our results support this finding as wild type cells treated with the Arp2/3 inhibitor CK-666 showed an orientation similar to the mutant, R196H. The effect of jasplakinolide on the wild type filament orientation might also be associated with an impaired Arp2/3 binding in the presence of jasplakinolide (37). How the defect in Arp2/3 binding contributes to the phenotype observed in patients needs to be determined in other cell types, such as neurons. However, the role of Arp2/3 was shown in regulating neural interconnectivity, through participation in functional maturation of dendritic spines (38). As abnormal spine structure has been shown in neurodevelopmental disorders associated with intellectual disability, the impaired interaction of the mutant actin with the Arp2/3 complex might play a role in the development of BWCFF.

Reduced cell stiffness as a result of the mutation is likely due to the reduced amount of F-actin detected in the mutant cells. The observed difference in the directionality of the fibers (fiber alignment) probably has a smaller contribution to the observed stiffness changes (26). However, we did not find any differences in the elasticity of wild type and mutant cells by applying a lower force regime by the DLOT. Based on the DLOT analysis the R196H mutation affects the process of force-driven tether formation. More specifically, within this step the observed reduction of the *F_peak_* and also the normalized *F*_peak_* values upon the mutation emphasize that wild type fibroblasts exhibit a higher mechanical resistance to the disruption of membrane-cytoskeleton adhesion in the mechanics of tether formation. Generation of the tether involves the force required to overcome membrane-cytoskeleton adhesion and viscous drag to allow cell protrusion. Despite the force being influenced by membrane mechanical properties like surface tension, curvature, and bending rigidity (39), no significant differences were found in membrane tension or bending modulus. This suggests that other factors, such as the interaction between the membrane and cytoskeleton or the efficiency of tether formation, may be more important in determining the force. The R196H mutation could impact these molecular interactions without affecting the membrane’s fundamental mechanical properties. Interestingly, other DLOT-derived parameters—such as effective tether viscosity, tether stiffness, and equilibrium tether force—were unaffected by the mutation, reinforcing the idea that the R196H-induced changes in cytoskeletal behavior are subtle and likely restricted to filament-membrane adhesion interfaces. The preserved bending modulus and membrane tension support the hypothesis that the primary structural alterations reside within the actin cortex and its interactions, rather than in the membrane itself. These data highlight the importance of local actin network architecture and binding strength over bulk mechanical properties in modulating cellular responses to external forces.

Cells continuously sense their environment and reorganize their actin cytoskeleton to withstand external forces (40). Actin remodeling is regulated by spatially and temporally controlled actin polymerization and depolymerization. The size and the density of F-actin network is regulated through the activation of actin-assembly factors by GTPase signaling cascades, F-actin barbed-end capping, and F-actin-disassembly factors (41). The availability of actin-ATP precursors does not limit actin polymerization, and polymerization can be induced in a regulated manner through the activation of Rho GTPases. Actin disassembly can also be promoted by several factors, e.g., the ADF/cofilin family, represented mostly by cofilin-1 in non-muscle cells. We found that the mutation accelerates cofilin dissociation from peripheral actin, but only during the response to the applied external force (uniaxial stretch). The question arises how the mutation might perturb the association of cofilin with actin. It is known that cofilin binds preferably to actin-ADP filaments, but it has also been proposed that cofilin binding to actin might accelerate Pi release. One possibility of how the mutation might affect cofilin binding is that the amino acid change in position 196 might allosterically act through the interaction network of the side chains of the backdoor of actin (Figure 1B), where P_i_ is leaving after ATP hydrolysis, since the binding site of cofilin is far away from the location of the mutated residue (Figure 6J). Another possibility for an allosteric regulation might be related to the mechanism how cofilin severs actin. The structural details of cofilin severing were revealed recently by Huehn and coworkers (42). Cofilin binds to two neighboring actin subunits (Figure 6J, subunit i and i-2) within the protofilament and keeps them together, while it changes the conformation of that subunit within the protofilament which is closer to the pointed end (Figure 6J). This results in the weakening of the contact between subunit i (Figure 6J, cyan) and the adjacent one (subunit i+2, red, not bound to cofilin) closer to the pointed end. This weakening is reflected in the disordered appearance of the D-loop in subunit i. By comparing the existing structural information for cofilin-bound actin, we analyzed the length of the possible salt bridges between the residues R196 and either E237 or E253 within the same actin protomer (Table 1). Apparently, these contacts are not affected by cofilin binding. However, comparison of the lateral contacts between the actin filaments pointed at the importance of the K113-E195 salt bridge in the weakening of interfilament contacts upon cofilin binding (Table 1). When only one cofilin molecule is bound to the actin filament, weakening of this salt bridge can be observed between the actin subunits i and i-1(Figure 6J). However, when all actin monomers are bridged by a cofilin molecule, the salt bridge between each protomer is weakened. In addition, another salt bridge can also be affected between R39 and E270, which was shown to be another important lateral contact stabilizing the filament (Figure 6K, Table 1) in the absence of cofilin. It is important to note that Pi release does not weaken the K113-E195 salt bridge, which, based on our analysis, is weakened by cofilin binding. Thus, based on these assumptions, cofilin binding might allosterically weaken the interstrand (lateral) contact, close to the residue R196 and possibly not via the backdoor of Pi release. Thus, conceivably, if this salt bridge is already perturbed by the mutation, then cofilin binding might be allosterically affected. Our hypothesis is that the mutation of the residue R196 might predispose the filament for breakdown through weakening of the lateral contact between its neighbouring residue E195 and the residue form the adjacent protomer, K113. Since this interaction is also sensitive to cofilin binding, this might influence the reorganization of actin by cofilin.

**Table 1.** Comparison of specific contacts within the actin protomer as well as the lateral contacts between the filaments and the effect of cofilin binding on them.

| PDB code (reference) | Actin | phalloidin/JASP/ cofilin | Interaction between residues |  | Distance (Angstrom) |
| --- | --- | --- | --- | --- | --- |
| Contacts within the actin protomer |  |  |  |  |  |
| 8DNH (67) | cytoplasmic, beta, Chains A-D | none | B:R195*:NH2 | B:E236*:OE1 | 4.03 |
|  |  |  | B:R195H*:NE | B:E252:OE2 | 3.30 |
| 6VAU (42) | rabbit muscle, Chains A-E | none | B:R196:NH1 | B:E237:OE2 | 3.76 |
|  |  |  | B:R196:NE | B:E253:OE2 | 3.47 |
| 6UBY (42) | rabbit muscle, Chains A-H | cofilin, chain I | E:R196:NH1 | E:E237:OE2 | 3.76 |
|  |  |  | E:R196:NE | E:E253:OE2 | 3.47 |
| 6VAO (42) | rabbit muscle, Chains A-E | cofilin, chains J-I | D:R196:NH2 | D:E237:OE2 | 4.47 |
|  |  |  | D:R196:NE | D:E253:OE2 | 2.84 |
| 5YU8 (68) | chicken muscle, Chains A-E | cofilin, chains H-J | D:R196:NH1 | D:E237:OE2 | 4.87 |
|  |  |  | D:R196:NE | D:E253:OE1 | 2.95 |
| Lateral contacts between the filaments |  |  |  |  |  |
| 8DNH (67) | cytoplasmic, beta, Chains A-D | none | A:K112*:NZ | B:E194*:OE2 | 3.62 |
|  |  |  | A:E269*:OE1 | B:R38*:NH1 | 2.39 |
| 6VAU (42) | rabbit muscle, Chains A-E | none | A:K113:NZ | B:E195:OE1 | 4.94 |
|  |  |  | A:E270:OE1 | B:R39:NH2 | 3.56 |
| 8OI6 (23) | cytoplasmic, beta, Chains A-D | phalloidin, H-J | A:K113:NZ | B:E195:OE1 | 4.51 |
|  |  |  | A:E270:OE1 | B:R39:NE | 3.31 |
| 6T23 (69) | rabbit muscle, Chains A-E | JASP | D:K113:NZ | C:E195:OE1 | 2.88 |
|  |  |  | D:E270:OE2 | C:R39:NH1 | 2.79 |
| 6T24 (69) | rabbit muscle, Chains A-E | JASP | D:K113:NZ | C:E195:OE1 | 3.25 |
|  |  |  | D:E270:OE2 | C:R39:NH1 | 2.76 |
| 6UBY (42) | rabbit muscle, Chains A-H | cofilin, chain I | G:K113:NZ | E:E195:OE1 | 8.42 |
|  |  |  | G:E270:OE1 | E:R39:NH2 | 3.49 |
| 6VAO (42) | rabbit muscle, Chains A-E | cofilin, chains J-I | A:K113:NZ | D:E195:OE2 | 10.79 |
|  |  |  | A:E270:OE1 | D:R39:NH2 | 7.79 |
| 5YU8 (68) | chicken muscle, Chains A-E | cofilin, chains H-J | E: K113:NZ | D:E195:OE2 | 10.78 |
|  |  |  | E: 270:OE1 | D:R39:NH2 | 5.20 |

Cofilin is ubiquitous and abundant in neurons and plays a crucial role in neurofilament dynamics through its effects on dendritic spines and the trafficking of glutamate receptors, a mediator of excitatory synaptic transmission (43). Regulation of cofilin involves various signaling pathways converging on LIM domain kinases (LIMK1/2) and slingshot phosphatases (SSH1/SSH2), which phosphorylate/inactivate and dephosphorylate/activate cofilin, respectively. Abnormalities in cofilin regulation have been associated with several neurodegenerative disorders (44). Therefore, the question arises whether targeting cofilin (by inhibiting its phosphorylation to strengthen its interaction with actin) might provide a potential to treat BWCFF-mutation-related brain disorders, such as epilepsy.

Taken together, we found that the BWCFF-mutation, R196H causes a decrease in the amount of actin filament bundles in patient-derived fibroblasts. This might be related to the reduced stiffness of the mutant cells determined by AFM. In addition, we found that in the presence of the mutation two actin-regulating proteins, the Arp2/3 complex and cofilin show impaired binding to the actin filament, the latter is important during actin reorganization. Based on these findings the question arises which of these changes affect the phenotype of the patients and whether that defect can be rescued biochemically. To answer these questions, we have to investigate the effect of this mutation in the clinically relevant context, e.g. affected cells, such as neurons and provide strategies to rescue the defects in polymerization or in binding to either the Arp2/3 complex or cofilin. Dendritic spine density in neurons correlates with intellectual ability (45). The dendritic actin cytoskeleton is composed of two types of structures: a stable, cortical actin ring providing mechanical support and dynamic actin filaments necessary for synaptic plasticity (46). Cofilin activity is instrumental for the dynamic plasticity of dendritic spines, and this is critical in normal physiological conditions, such as learning. Thus, dysregulation of F-actin stability or cofilin activity might contribute to the structural and dynamic plasticity of dendritic spines (47). Proper Arp2/3 binding to actin has a distinct role, as it is dispensable for the emergence of dendritic filopodia, but it is indispensable for their functional maturation to dendritic spines during development (38).

## MATERIALS AND METHODS

### Primary antibodies for western blotting (WB)

β-actin antibody, Cat# MCA5775GA from Bio-Rad; pan-Actin antibody, clone 2A3 Cat# MABT1333 from Sigma Aldrich; GAPDH antibody, Cat# 97166 from Cell Signaling Technology, tubulin antibody, Cat# PA5-58711, from Thermo Scientific.

### Primary antibodies for immunofluorescence (IF)

cofilin, Cat# ab42824 from Abcam.

### Secondary antibodies for WB

Peroxidase AffiniPure Goat Anti-Mouse IgG (H+L), Cat# 115-035-003; Peroxidase AffiniPure Goat Anti-Rabbit IgG (H+L), Cat# 111-035-003, both from Jackson ImmunoResearch.

### Secondary antibodies for IF

Chicken anti-Rabbit IgG (H+L) Cross-Adsorbed Secondary Antibody, Alexa Fluor 488, Cat# A21441; Alexa Fluor™ Plus 647 Phalloidin, Cat# A30107; both from ThermoFisher Scientific. Abberior star 635P conjugated to phalloidin for STED imaging, Cat# ST635P-0100-20UG from Abberior.

### Live cell stain

CellMask™ Orange Actin Tracking Stain, Cat# A57247 from Thermo Scientific.

### Cell culture and cell treatments

Patient-derived wild type and mutant fibroblast cells were isolated as described in (48) and were cultured in BioAFM2 medium. All cell culture was performed at 37°C in a humidified atmosphere containing 5% CO_2_. For specific experiments cells were treated with 0.1 μM jasplakinolide (Cat# 420127, Merck) for 30 min or an Arp2/3 inhibitor, CK-666 (Cat# SML0006, Merck) was applied in 100 μM concentration for 60 min at 37°C. Wild type cells were treated with 5 μM latrunculin A (Cat# 428021, Merck) for 20 minutes and processed for further DLOT analysis.

### Determination of F-actin and G-actin content

Samples of wild type (#60051) and R196H mutant (#1620) fibroblasts were pre-treated with 0.1 μM jasplakinolide (Cat# 420127, Merck) for 30 minutes (or with DMSO used as a control) prior to lysis. Lysis was carried out according to the protocol described in the G-actin / F-actin In Vivo Assay Kit (Cat# BK-037, Cytoskeleton Inc.). After a preclearing centrifugation step, G-actin and F-actin were separated by ultracentrifugation. The supernatant containing G-actin, and the pellet (resuspended in milliQ water) containing F-actin was loaded on an SDS-PAGE gel and the amount of actin was visualized by using a β-actin or a pan-actin antibody. After HRP-conjugated secondary antibody incubation, the membranes were incubated with chemiluminescence substrate and developed on Hyperfilms. Bands were quantified using ImageJ 1.53c.

### Immunoblotting of cell lysates

Wild type (#60051) and R196H mutant (#1620) fibroblasts were harvested in 25 mM HEPES, pH 7.4, 150 mM NaCl, 1 mM EGTA, 1% NP-40, 10% glycerol, supplemented with the following protease and phosphatase inhibitors: 10 mM sodium pyrophosphate, 10 mM sodium fluoride, 5 mM sodium vanadate, 1 mM PMSF and cOmplete, EDTA-free protease inhibitor cocktail (Sigma, Cat# 4693132001). Lysates were centrifuged with 5000 rpm for 5 minutes at 4°C and the supernatant was snap frozen for further immunoblotting. Proteins were separated using standard SDS-PAGE gel electrophoresis and immunoblotted as described above.

### Monolayer stretching

Fibroblast monolayers (wild type, #60051 and #52556; R196H, #1620 and #49113) were cultured on special chambers (CuriBio Inc., Cat #CS-2×25-UPF) used in the Cytostretcher LV instrument of CuriBio. The bottom of the chamber was treated with 0.2 mg/ml polydopamine solution (dissolved in 10 mM TRIS buffer, pH 8.5) for 2.5 hours, washed 3 times with sterile water to create a hydrophilic surface for gelatin coating. Stretching was carried out with 0.5%/sec velocity and the monolayer was kept stretched for 15 minutes before fixation. 2‧10^4^ cells were seeded in 5 mm x 5 mm chambers and were grown for 36 hours before stretch.

### Immunofluorescence staining of fixed monolayers

Cells grown on cytostretcher chambers (either non-stretched or stretched) were fixed with Image-iT™ Fixative Solution (ThermoFisher Scientific, Cat# R37814) for 15 minutes. After that, cells were washed with HBSS, permeabilized (0.25% Triton X-100 in TBS-T, 10 min RT), blocked (1% BSA in TBS-T, 1 hour RT), and incubated with the primary antibody (dilutions prepared in 1% BSA-TBS-T for cofilin – 1:200; incubation was done overnight at 4°C). After thorough washing in TBST, cells were stained simultaneously with the appropriate secondary antibodies (dilutions were prepared as 1:1000) and phalloidin (1:1000 in 1% BSA-TBST) for 1 hour at RT, washed in TBS-T and PBS prior to imaging.

### Confocal and STED microscopy

Non-stretched and stretched samples were analyzed on a Nikon Ti2 confocal microscope. The field of view for imaging was a 140 µm × 140 µm area (resolution: 1000 × 1000 pixels) and pictures were taken by using a 20× lens (numerical aperture: 0.75). Live cell images (wild type, #60051; R196H, #1620) were taken by using a 60x lens (numerical aperture: 1.40, oil). The field of view for imaging was a 120 µm × 120 µm area (resolution: 1000 × 1000 pixels). STED images of fixed cells (wild type, #60051; R196H, #1620) were taken by using a 100x lens (numerical aperture: 1.45, oil). Depletion was carried out with a 775 nm laser, with 20% laser power. Images were taken from randomly selected areas of the cell monolayer.

### Quantification of actin filament bundle width from the STED images

STED images were deconvoluted by using the Huygens Professional software 23.04, Scientific Volume Imaging BV. Quantification of actin width was carried out after the deconvolution by using a macro plugin of ImageJ 1.53c, which is using a Gaussian fit of the line profile of phalloidin fluorescence. The width is given as the fitted value corresponding to the 2x standard deviation (2xSD) of the Gaussian distribution. All analyses were performed blinded, such as data analyzers were unaware of the genotypes and treatments.

### Quantification of the directionality profile of the actin filament bundles

Directionality analysis was carried out using the directionality plugin for ImageJ 1.53c. The analysis of an image (120 µm × 120 µm area) results in a histogram reflecting the number of structures in the image that are subjected to a certain angular direction. The program fits a gaussian curve. The “amount” value, derived from the mean value of the Gaussian fit, indicates the percentage of the present structures that align in the specific direction of the highest peak. The standard deviation (SD) of the Gaussian fit corresponds to the value of dispersion. All analyses were performed blinded, such as data analyzers were unaware of the genotypes and treatments.

### Quantification of cofilin relocalization upon stretch

First, confocal images measured in the z direction were summed by using the ImageJ 1.53c “Maximal Intensity Projection” function. After that, stacked images of both phalloidin- and cofilin-stained samples were converted to a binary image by using a threshold between 80-90% to identify the areas having a fluorescence signal. Then, the Particle analysis tool of ImageJ 1.53c was applied on the masked images (to identify the areas having “no” fluorescence signal and a size limitation of 20-100 μm^2^ was used to exclude tiny and very large holes) to determine the size of each “empty” area (Supplementary Figure 3). After that, the corresponding “empty” areas of phalloidin- and cofilin-stained images were used for further calculations. We subtracted the area of the phalloidin-stained “holes” form the area of the cofilin-stained “holes” and divided by the area of the phalloidin-stained “holes” to get the ratio of the area where cofilin staining cannot be detected with the used threshold. Then, the numbers were converted to percentage for each condition (wild type or mutant, stretched or non-stretched) were plotted and compared. All analyses were performed blinded, such as data analyzers were unaware of the genotypes and treatments.

### Preparation of recombinant actin and sample preparation for AFM imaging

Recombinant actin was produced and purified as described in (21). G-actin was polymerized overnight at 10 µM concentration in a polymerization buffer containing 100 mM KCl, 2 mM MgCl_2_, and 0.05 mM EGTA. Actin filaments, stabilized with phalloidin (Cat# P1951, Merck) at a 1:1 molar ratio, were introduced to the lipid bilayer at 2 µM concentration. After a 15-minute incubation period, actin paracrystals were successfully formed on the bilayer surface.

### AFM imaging of recombinant actin

Lipid vesicle preparation and lipid bilayer formation on Mica: A 1:1 molar ratio mixture of DPPC (Dipalmitoylphosphatidylcholine) and DPEPC (1,2-dioleoyl-sn-glycero-3-ethylphosphocholine) was hydrated in a buffer solution consisting of 10 mM Tris, 100 mM NaCl, and 3 mM CaCl_2_ at pH 7.4, to achieve a final lipid concentration of 1 mM. The lipid vesicles were subsequently formed through extrusion using a 100 nm polycarbonate membrane. A freshly cleaved mica surface was treated with 100 µL of the lipid mixture and incubated at room temperature for 30 minutes. Following this incubation, the temperature was elevated to 55 °C for 15 minutes to induce the rupture of vesicles, thereby forming a positively charged lipid bilayer. The structural and topological features of the actin filaments were examined using a Cypher Atomic Force Microscope (Asylum Research, Santa Barbara, CA) in non-contact mode. Silicon cantilevers (BL-AC40TS, Olympus, Tokyo, Japan) with a nominal tip radius of 8 nm and a resonance frequency of approximately 25 kHz were employed. Imaging was performed in a hydrated environment to preserve the native conformation of the lipid bilayer and the actin filaments.

### AFM force spectroscopy of patient-derived fibroblasts

AFM imaging (Igor Pro 6.37 software) was performed in contact mode with an MFP3D AFM (49) using Bruker, MSCT-A probes (nominal typical spring constant = 70 pN/nm). Cantilevers were calibrated by the thermal method (50). Cell monolayer samples (wild type, #60051 and #52556; R196H, #1620) were grown on circular microscope slides, which were mounted in the Bio-Heater module of the AFM. Imaging and force spectroscopy were carried out in a temperature-controlled liquid environment at 37°C. First an AFM image was taken, then *in situ* force spectroscopy was carried out collecting 100 force curves on selected 3×3 µm regions of cell surfaces (force mapping). The individual force curves were recorded with a vertical Z-piezo movement speed of 1 μm/s, until the force set point (0.5 nN) was reached, then the tip was retracted. Elastic moduli were obtained by fitting the indentation curves with the blunted pyramidal model as described in Rico *et al*. (51). For the calculations (52) the irregular pyramid shape with a semi-included angle of 20°, and with a spherical cap radius of 10 nm was used as described for the Bruker, MSCT-A probe. Poisson ratios of the tip and the sample were set to 0.2 and 0.5, respectively.

### Mechanical protocol for indentation and tether formation in a DLOT system and data analysis

The mechanical protocol used to study the viscoelastic properties of the cytoskeleton-membrane system with the DLOT consisted of a push-and-hold and ramp-and-hold protocol under position feedback (red trace in Figure 5B). A laser-trapped bead (3 µm in diameter, NH_2_-functionalized from Kisker Biotech GmbH & Co. KG, Cat# PPS-3.0NH2P) was held in a fixed position near a cell (wild type, #52555 or R196H mutant, #49113 fibroblast) firmly anchored to the bottom of the experimental chamber. The cell was moved along the x-y plane by the nano-positioner at the velocity of 1-4 µm/s during both the indentation phase and the tether formation. The velocity range was chosen to be in the permeation regime (1–100 µm/s) (53), where transmembrane proteins remain anchored to the cytoskeleton while membrane lipids flow around them. This dynamic interaction is responsible for the friction forces between the lipid bilayer and the cytoskeleton that explains the rise of membrane tension. The mechanical perturbation exerted on the cell were applied approximately 5 µm away from the nucleus to make negligible the contributions from the stiffer nuclear region, which could otherwise dominate the overall mechanical response during membrane deformation due to its stiffness 5–10 times higher than the surrounding cytoskeleton (54).

The force response to the mechanical protocol can be divided into three main phases: indentation, tether formation and elongation, return to the “initial position”. The indentation can be further divided into three phases (Figure 5B). [1] Approach (white area): the cell is moved toward the bead at constant velocity (2 µm/s, Figure 5B) until contact occurs. [2] Elastic Indentation (light green): a compressive (negative) force develops with indentation depth, allowing a Young’s modulus to be determined using the Hertz model (55). The phase concludes at the end of the ramp, where the force reaches a value *F_i_*. (Enlarged in Supplementary Figure 2A) [3] Relaxation (light violet): following the end of the ramp, the compressive force decreases during the hold period, reaching a nearly steady force F_r_ (Supplementary Figure 2A). The tether formation and elongation also consist of three different processes. [4] Tether Initiation (yellow): ramp-shaped pull at 4 µm/s induces an early force peak resisting to extension (positive) (*F_peak_*), followed by a sharp force drop, indicating the weakening of the interaction between the plasma membrane and the cytoskeleton (56). [5] Tether Elongation (pink): positive force increases again with further pulling and tether extension, reaching *F_1_* at the ramp’s end. Ramps with different lengthening, and consequent tether elongation, of 10, 20, 30, 40 and 50 μm were applied. [6] Tether Relaxation (light blue): during the hold phase following the end of pull, the force relaxes to an equilibrium value (*F_e_*). The last phase is covered by two processes. [7] Tether retraction (light gray): the cell is returned to its initial position using the same velocity as in the pulling phase, and [8] Hold time (dark gray): a 15-second hold, after which a new cycle of tether elongation/retraction begins.

For the second cycle there is no need for preliminary contact and indentation because the tether pre-exists from the first cycle, thus the cycle starts from the tether elongation (phase 5’). The pre-existing tether bypasses also the *F*_peak_ response characterizing membrane-cytoskeleton interaction. Thus, for the subsequent pulling cycles, no *F*_peak_ is observed as the tether remains connected. Notably, also by direct inspection it is evident that the relaxation force response *F(t)* during the hold periods following indentation (phase 3) and tether formation (phase 6) have a biphasic time course with a faster and a slower component.

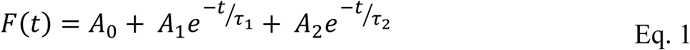

This behaviour suggests an equivalent mechanical model with five-elements (Zener model) (Supplementary Figure 2B) (57, 58). From the values of the amplitudes *A*_0_, *A*_1_, *A*_2_ and relaxation times τ_1,_ τ_2_, the model enables the estimation of both the undamped (*k*₀) and damped elastic coefficients (*k*₁ and *k*₂), as well as the friction coefficients (γ₁ and γ₂), in the two different processes (holds following indentation and tether elongation). The shorter τ corresponds to the Marangoni convective flow of lipids from regions of low membrane tension to regions of high membrane tension, while the longer τ is associated with the diffusive flow of lipids (57, 59–61).

### Mechanical characterization of cell membrane indentation using DLOT and calculation of the elastic modulus

Indentation response (phases 1-3 of DLOT, Figure 5B) of wild type fibroblast cell membrane to a ramp-shaped push (***v*** = 4 μm/s) is illustrated in Supplementary Figure 2A. The start of the indentation is marked by the time (indicated in the Figure by the magenta vertical dashed line) at which the position of the membrane in the contact point (*L* trace, black) starts to deviate from the position imposed by the nanopositioner (*X* trace, red), and indentation force develops (*F*, blue, as measured by the displacement of the bead from the trap center multiplied by the trap stiffness). The indentation depth (*d*) is measured at any time after the contact by the difference (*X* – *L*). During the ramp, when applied over a short duration (*v* ≥ 4 µm/s), both force and indentation depth increase linearly up to a value attained at the end of the ramp, *F*_i_ and d_i_ respectively, suggesting an elastic response. Following the ramp-end, there is a force relaxation, which implies the return of the bead toward the trap center with further movement of the membrane and corresponding further increase in indentation. This phase reveals the viscoelastic nature of the response to indentation that reaches a near-equilibrium state characterized by a force *F*_r_ and an indentation *d*_r_. Both force and indentation relaxations are fitted by a double-exponential decay equation (orange dashed and yellow dashed lines respectively). The biexponential process is the same as the two signals are coupled by the constant trap compliance.

To estimate the relevant mechanical parameters of the viscoelastic response with the equivalent mechanical model (Zener model, Supplementary Figure 2B) would require that force relaxation following the ramp end, occurred without further changes in *L* and thus *d*, a condition that could be achieved only neutralizing the trap compliance under length clamp conditions (62). In the presence of the trap compliance, the estimate of the viscoelastic parameters derived from the relaxation process must be considered an approximation and only the “static” elastic constant (*k*_0_), characterizing the steady state response (achieved at sufficiently long times after the end of the push) is quantitatively correct. Consequently, the mechanical model in Supplementary Figure 2B is simplified, and only account for *F*_i_ and *F*_r_. For *F*_i_ the response is attributed to the parallel of the three springs *k*_0_+*k*_1_+*k*_2_ and for *F*_r_ only to the static elastic component *k*_0_ at the end of relaxation. In this way two apparent elastic moduli can be estimated through equation 2 (Sneddon–Hertz model): the “instantaneous” elastic modulus *E*_i_, given by the parallel of the three springs, and the “static” elastic modulus *E*_r_, due to the only element *k*_0_.

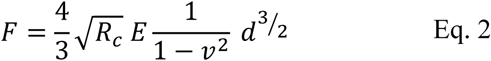

Where *R*_c_ represents the radius of the bead (1.5 µm), *E* the apparent Young’s modulus (either *E*_i_ or *E*_r_), *v* the Poisson ratio, that was assumed 0.5, and *d* the indentation depth. The Poisson’s ratio is defined as:

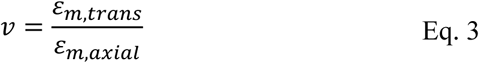

where:

ε_m,trans_ is transverse strain (negative for axial traction, positive for axial compression)

ε_m,axial_ is axial strain (positive for axial traction, negative for axial compression). From Eq. 2:

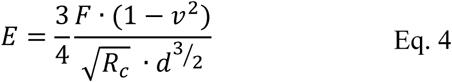

### Measurement of the membrane tether radius

The radii of membrane tethers were estimated using bright-field imaging on the DLOT system. After tether force reached equilibrium, bright-field images were acquired using the DLOT setup (wavelength: 450 nm, 60X water immersion objective, NA 1.2, exposure time for single images 50 ms). Calibration of the transverse (x and y) planes was performed by displacing a micropipette via the nanopositioner, covering known distances from 0.5 to 20 μm in both directions.

The image shadow was subtracted, and background noise minimized using ImageJ/FIJI software (version 2.9.0, National Institute of Health, Madison, WI, USA) through a sequence of image processing steps. These included conversion to grayscale, background subtraction using the “subtract background” function, thresholding, creation of a binary mask, the Despeckle tool, and averaging 100 frames per tether via the Z-project method. Measurements were restricted to sharply focused sections of the tethers to avoid errors from shadowing effects, as membrane tethers are often angled relative to the coverslip.

The resolution of the systems is half the diameter of the Airy disk on the detector, calculated as 1.22 λ/NA. The intensity distribution of the Airy disk across its centre can be approximated by a gaussian function with width σA. This was measured using 200 nm fluorescent beads (TetraSpeck microspheres; Invitrogen) which are small enough to be considered point-like sources, yet large enough to generate a strong and measurable signal (63). By fitting the observed profile of the tether with a gaussian width σ_O_ and deconvolving we can derive the width σ_t_ of the Gaussian profile that best fits the tether deconvolved image:

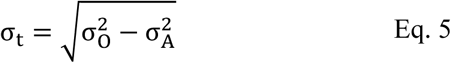

The radius of the tether is assumed to correspond to r_t_ ≈ 2 σ_t_, when the intensity of the gaussian has reduced to ≈ 5% the peak intensity.

### Calculation of the membrane tension and the bending modulus

The membrane tension, *T* for an equilibrium force *F*_e_ can be estimated with the following equation (29):

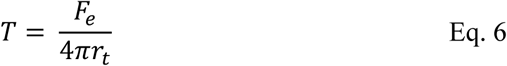

where *r*_t_ is the tether radius.

The energy required to deform a cell’s membrane depends on the elastic resistance opposed to the bending of the membrane, expressed by the bending modulus (*e*_b_). *e*_b_ for a tether with the equilibrium force *F*_e_ can be calculated using Eq. 7 (64):

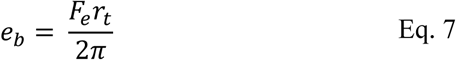

where *r*_t_ is the tether radius. A higher *e*_b_ indicates a larger resistance to membrane deformation. Biological membranes typically exhibit *e*_b_ values in the range of 10^−20^−10^−18^ J, depending on membrane composition and cytoskeletal interactions.

### Calculation of the normalized *F_peak_* (*F\**) values

Given that *F*_peak_ is not a single physical point but it arises over a circular contact area between the bead and the membrane with a radius *R*_p_ (radius of the region of the membrane that is locally deformed by a force that leads to formation of the tether), theoretical modelling accounting for the ratio of *R*_p_ to the tether radius (*r*_t_) is necessary to evaluate *F*_peak_ in relation to the equilibrium force (*F*_e_) (65).

The asymptotic relationship for *R*_p_ >> *r*_t_ is described by the equation:

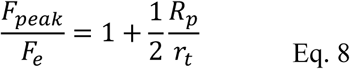

Applying this equation to our data, we could normalize *F*_peak_ for *R*_p_ obtaining *F\**_peak_ (pN/nm^2^), Eq. 9.

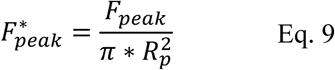

Calculation of *F\**_peak_ requires to know the tether radius *r*_t_ and then to estimate *R*_p_. The calculations were carried out using *F*_peak_ and *F*_e_ values measured at a constant tether extraction speed (4 µm/s) up to a constant length (L, 38-43 µm), and only on the traces for which *r*_t_ could be accurately measured by DLOT.

### Calculation of the tether effective viscosity (γ_eff_)

During tether elongation at a constant velocity (Figure 5B, phase 5), force rises reaching the value of *F_1_* at the end of the ramp. The rate of force rise decreases monotonically in an almost exponential way as expected from a viscoelastic response of the cell membrane. The resistance to elongation and thus *F_1_* depends on the velocity of elongation. During the hold phase (Figure 5B, phase 6) following the elongation, the force relaxes, displaying a biexponential decay pattern until it stabilizes at an equilibrium force value *F_e_*. For accurate determination of the elastic and viscous parameters, the ramp and hold times must be sufficiently long to allow the dashpot with the higher time constant (friction coefficient over elastic constant) to load and relax to steady state. Specifically, the extent of membrane elongation selected for determining *F_1_* must imply times larger than the force rise time *t_r_* (the time required for the force to increase from 10% to 90% of its maximum change). This is the condition for the damped elastic component to be strained to the force corresponding to the viscous load for the imposed lengthening velocity. This condition is satisfied for a ramp rate of 4 µm/s when the ramp duration exceeds 6 seconds, given that 5 < *t_r_* < 7 s. Consequently, it applies to ramps longer than 28 µm. Therefore, only ramps with L > 35 µm were used to determine the tether’s effective viscosity.

### Statistical Analyses

Statistical analyses were carried out in GraphPad Prism 4.01. Significance was determined by one-way or two-way ANOVA followed by a Bonferroni post-test. Differences between groups were considered statistically significant if *p* < 0.05.

## AUTHOR CONTRIBUTIONS

É.G., E.B., Á.A, P.B. and A.V. designed the experiments. É.G., E.B., I.P., T.B., Á.A, L.H., P.B. and A.V. performed experiments. É.G., E.B., Á.A, K.P., L.H., T.B. and A.V. analyzed data. É.G., Á.A., K.P., JN.G., I.P., M.R., N. DD., K.M., P.B. and A.V. wrote the manuscript. All authors contributed feedback for the manuscript. All authors have read and agreed to the published version of the manuscript.

## ACKNOWLEDGEMENT

The authors thank the members of the PredACTINg Consortium for helpful discussions. The technical contribution of Béláné Szénási from the Department of Molecular Biology of Semmelweis University with ultracentrifugation is greatly acknowledged. The authors also thank Dr. Zsolt Mártonfalvi for the introduction of specific functions of Igor Pro 6.37.

## DATA AVAILABILITY STATEMENT

The original contributions presented in the study are included in the article. Further inquiries can be directed to the corresponding authors.

---

**Supplementary Figure 1, related to Figure 3.**
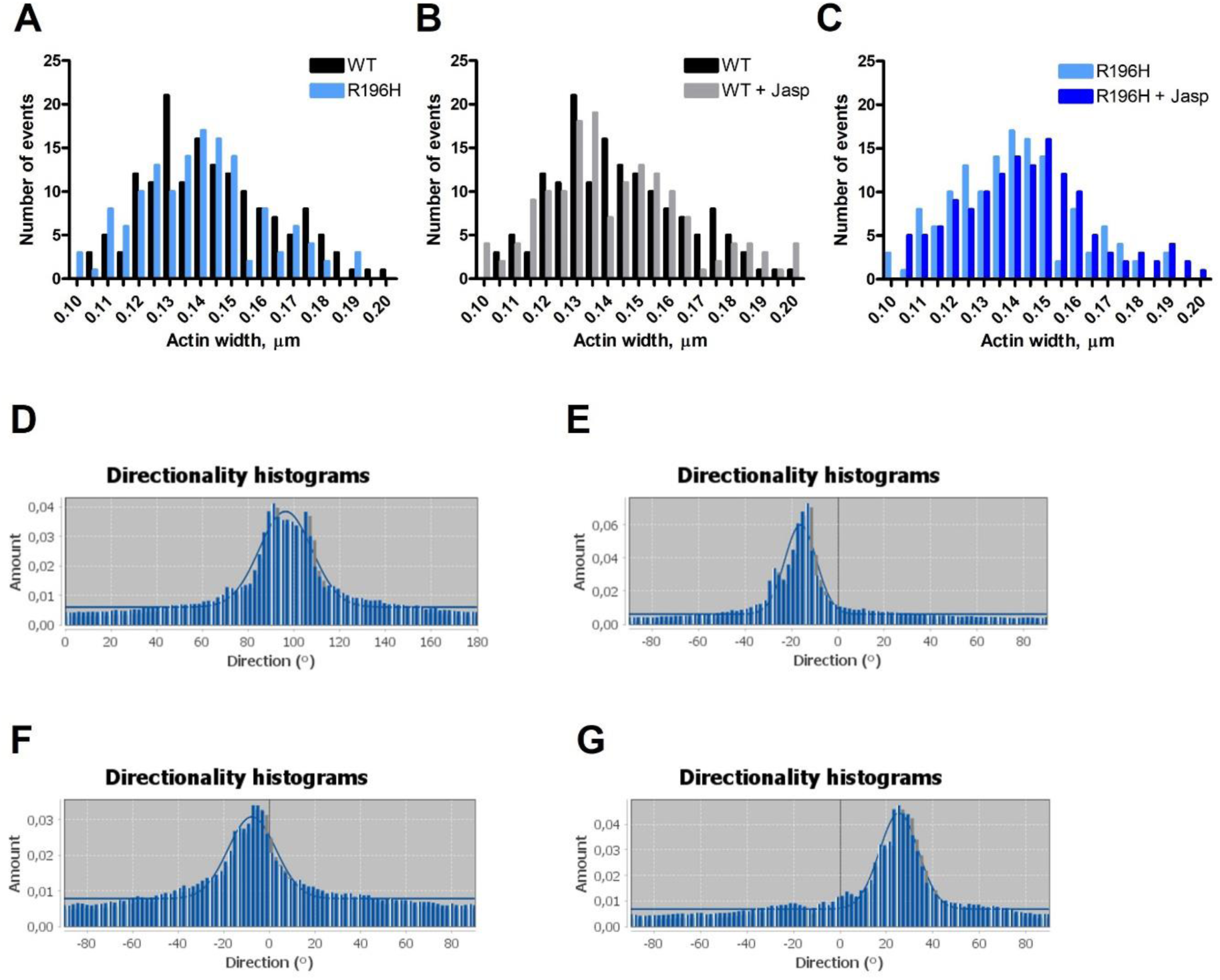
Pairwise comparison of the histograms of actin width determined for wild type and the R196H fibroblasts **(A)**, or to compare the effects of jasplakinolide either on wild type **(B)** or on the R196H fibroblasts **(C)**. Examples of directionality histograms created by ImageJ, 1.53c for the pairs of wild type **(D)** and the R196H **(E)** fibroblasts related to Figure 3C-E, as well as for the pairs of wild type fibroblasts treated with either DMSO **(F)** or CK-666 **(G)** related to Figure 3F and G.

**Supplementary Figure 2, related to Figure 5.**
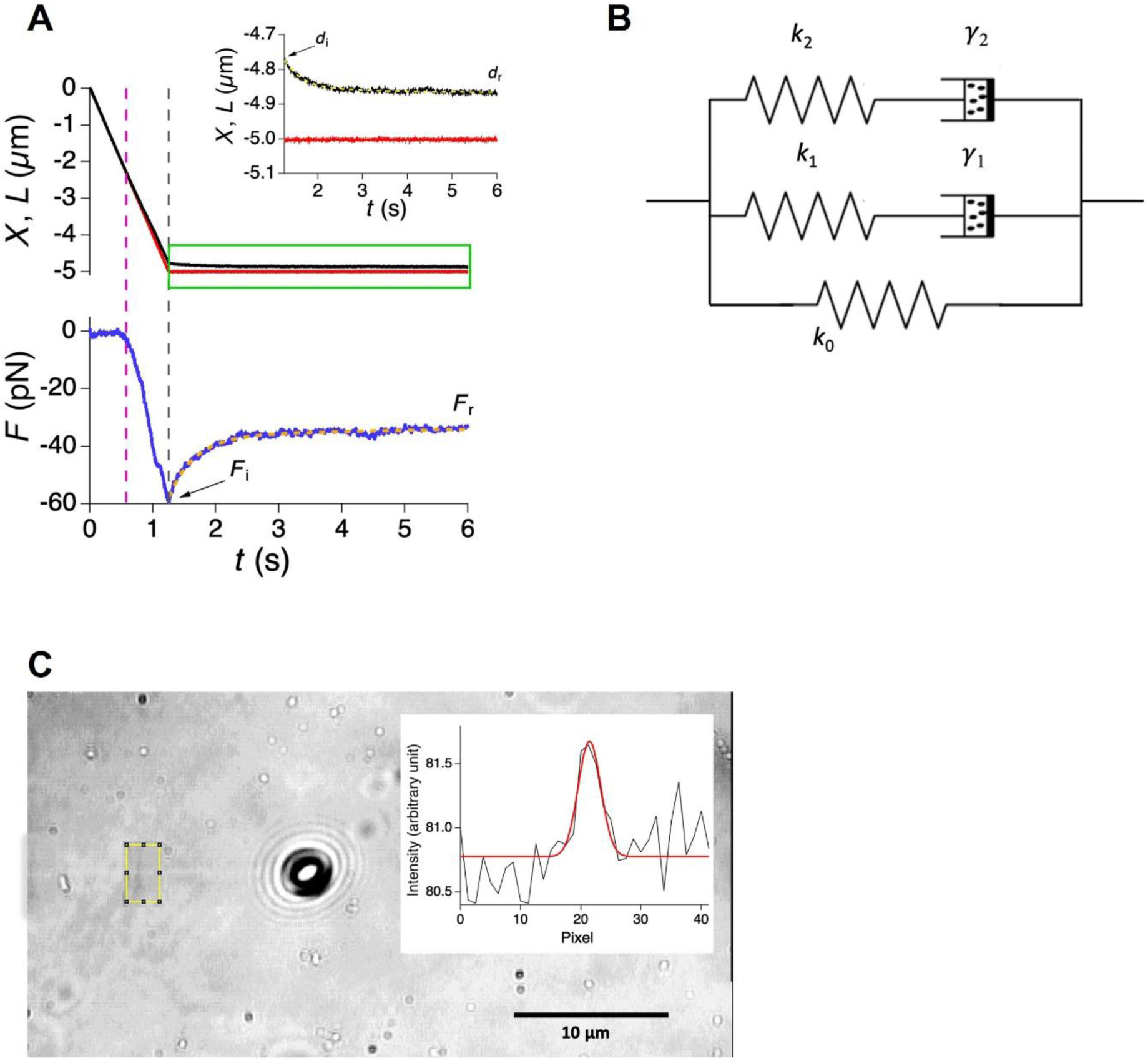
**(A)** Cell membrane indentation with a ramp-shaped push (*v* = 4 μm/s). The indentation begins when the membrane’s position (*L*, black) deviates from the nanopositioner (*X*, red), indicating the development of resisting force (*F*, blue). Force increases linearly during the ramp, followed by a relaxation phase, revealing the viscoelastic nature of the response. The relaxation follows a double-exponential decay and reaches near equilibrium at *F*_r_. Inset shows vertically zoomed *L* and *X* traces within the green box. *d*_i_ and *d*_r_ show the indentation depth at the end of the push and after the relaxation, respectively. Orange and yellow dashed lines: double-exponential fit on the traces. The magenta vertical dashed line marks the start of the indentation, the black vertical dashed lines indicate the ramp ends. **(B)** Schematic representation of a five-element Zener model used for identification of the mechanical parameters explaining the response of the system. The model in its simplest version (Maxwell model) comprises two parallel branches: one with a spring (*k_0_*) representing the purely elastic component (static), and the other with a series combination of a spring (*k_1_*) and dashpots (γ*_1_*) representing the viscoelasticity responsible for the relaxation process. The biphasic appearance of the relaxation can be accounted for by adding in parallel a second series combination of a spring (*k_2_*) and a dashpot (γ*_2_*) (Zener model). **(C)** Tether size measurements using the DLOT imaging. The figure shows a representative image of a tether extracted from wild type fibroblasts. Inset illustrates the transversal intensity profiles of the tether measured within the yellow dashed rectangles. To compare the tether radius among different experimental models, avoiding possible changes associated with tether length, all measurements were conducted on tethers with a narrow range (38–43 μm) of lengths.

**Supplementary Figure 3, related to Figure 6.**
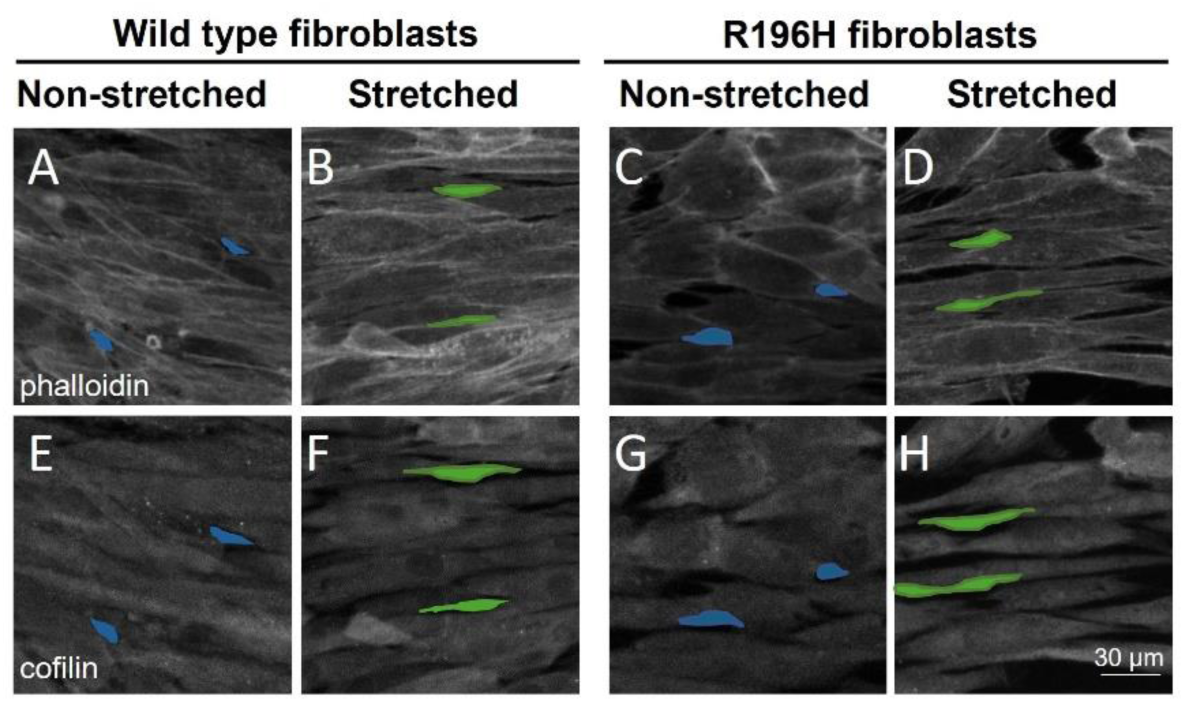
**(A-H)** Immunofluorescence analysis of wild type (A, B, E, F) or R196H (C, D, G, H) fibroblasts kept either non-stretched (A, C, E, G) or stretched (B, F, D, H) for 15 minutes. Samples were stained with either phalloidin (A-D) or for cofilin (E-H). Examples of areas detected in non-stretched (blue) or stretched (green) confocal images for visual comparison are shown.

**Supplementary Table 1.**
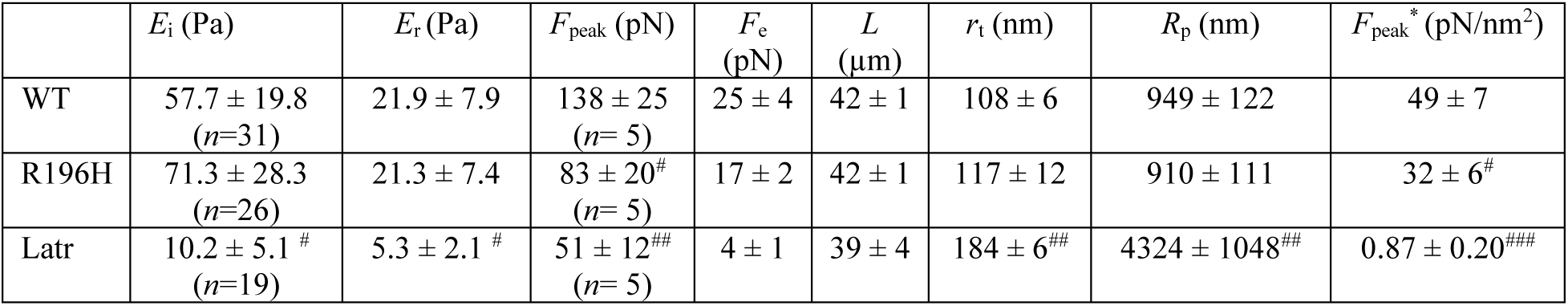
Mean values ± SD of apparent elastic modulus (*E*_i_) and equilibrium elastic modulus (*E*_r_), *F*_peak_, *F*_e_, tether length (*L*), tether radius (*r*_t_), patch radius (*R*_p_), and normalized peak force (*F*_peak_*) for all experimental conditions. Measurements were performed on wild type fibroblasts (WT), fibroblasts expressing the R196H mutation, and wild type fibroblasts treated with 5 μM Latrunculin A (Latr). *R*_p_ was calculated based on the tether radius (*r*_t_) measured using the DLOT imaging setup. *F*_peak_ and *F*_e_ values were analyzed at tether lengths ranging from 38 to 43 μm, with a retraction speed of 4 µm/s. *n* indicates the number of cells analyzed under each specific protocol: push or pull. Statistical significance vs. WT: # *p* < 0.05; ## *p* < 0.01; ### *p* < 0.001.

| | $E_i$ (Pa) | $E_r$ (Pa) | $F_{\text{peak}}$ (pN) | $F_e$ (pN) | $L$ ($\mu\text{m}$) | $r_t$ (nm) | $R_p$ (nm) | $F_{\text{peak}}^*$ (pN/nm<sup>2</sup>) |
| --- | --- | --- | --- | --- | --- | --- | --- | --- |
| WT | 57.7 ± 19.8<br>($n=31$) | 21.9 ± 7.9 | 138 ± 25<br>($n=5$) | 25 ± 4 | 42 ± 1 | 108 ± 6 | 949 ± 122 | 49 ± 7 |
| R196H | 71.3 ± 28.3<br>($n=26$) | 21.3 ± 7.4 | 83 ± 20<sup>#</sup><br>($n=5$) | 17 ± 2 | 42 ± 1 | 117 ± 12 | 910 ± 111 | 32 ± 6<sup>#</sup> |
| Latr | 10.2 ± 5.1<sup>#</sup><br>($n=19$) | 5.3 ± 2.1<sup>#</sup> | 51 ± 12<sup>##</sup><br>($n=5$) | 4 ± 1 | 39 ± 4 | 184 ± 6<sup>##</sup> | 4324 ± 1048<sup>##</sup> | 0.87 ± 0.20<sup>###</sup> |

